# Pupillometry and electroencephalography in the digit span task

**DOI:** 10.1101/2021.10.21.465288

**Authors:** Yuri G. Pavlov, Dauren Kasanov, Alexandra I. Kosachenko, Alexander I. Kotyusov, Niko A. Busch

**Affiliations:** Institute of Medical Psychology and Behavioral Neurobiology, University of Tübingen, Tübingen, Germany; Department of Psychology, Ural Federal University, Department of Psychology, Ekaterinburg, Russian Federation; Institute of Psychology, University of Münster, Germany

**Keywords:** cognitive load, EEG, ERP, working memory, photoplethysmography, heart rate

## Abstract

This dataset consists of raw 64-channel EEG, cardiovascular (electrocardiography and photoplethysmography), and pupillometry data from 86 human participants recorded during 4 minutes of eyes-closed resting and during performance of a classic working memory task – digit span task with serial recall. The participants either memorized or just listened to sequences of 5, 9, or 13 digits presented auditorily every 2 seconds. The dataset can be used for (1) developing algorithms for cognitive load discrimination and detection of cognitive overload; (2) studying neural (event-related potentials and brain oscillations) and peripheral (electrocardiography, photoplethysmography, and pupillometry) physiological signals during encoding and maintenance of each sequentially presented memory item; (3) correlating cognitive load and individual differences in working memory to neural and peripheral physiology, and studying the relationship between the physiological signals; (4) integration of the physiological findings with the vast knowledge coming from behavioral studies of verbal working memory in simple span paradigms. The data are shared in Brain Imaging Data Structure (BIDS) format and freely available on OpenNeuro (https://openneuro.org/datasets/ds003838).

## Background & Summary

Working memory (WM) is the ability to hold stimulus representations in an active state and to manipulate these representations over brief time intervals after the original stimuli have already disappeared. Although WM is an essential mnemonic and executive function that is involved in numerous everyday tasks, its capacity is strictly limited to only a handful of items (Constantinidis & Klingberg, 2016; Luck & Vogel, 2013). Individual differences in WM capacity are correlated with measures of fluid intelligence and other cognitive abilities and with psychiatric disorders such as schizophrenia (Fukuda et al., 2010; Johnson et al., 2013).

One of the oldest WM tasks is the digit span task, dating back to 1887 (Jacobs, 1887). This task involves the sequential encoding of a string of digits (akin to briefly memorizing a telephone number), their maintenance over a few seconds, and their retrieval, with memory load ranging from within to beyond an individual’s capacity limit. Paradigms such as the digit span task have been used to study the cognitive and neural mechanisms underlying WM, and the nature of WM capacity limitations. However, despite its long history and popularity among experimental and clinical psychologists – auditory digit span task is a part of Wechsler Adult Intelligence Scale –, data on the neural basis of sequential memory encoding and maintenance have not been made public. Specifically, there is no publicly available dataset including electrophysiological and peripheral physiological responses on an item-by-item basis in the digit span task under different memory load conditions. Such a dataset would provide an essential resource for connecting more than a century’s worth of experimental psychological data with research on the brain mechanisms underlying cognitive load and working memory.

Here, we describe a multidimensional dataset containing electroencephalography (EEG), electrocardiography (ECG), photoplethysmography (PPG), pupillometry, and behavioral response data collected from 86 participants during performance of digit span task. The data are suitable for studying oscillatory brain activity and event-related brain potentials in response to increasing WM load starting from one item (digit) up to thirteen items. Moreover, this dataset links cognitive load with peripheral nervous system responses such as pupil dilation, shown to be indirectly related to the locus coeruleus norepinephrine system activity (Joshi et al., 2016; Murphy et al., 2014); heart rate variability; as well as pulse wave amplitude, reflecting vasoconstriction that can be used as a surrogate of autonomic and cortical arousal.

Using an auditory version of the digit span task provides an advantage for studying neural mechanisms of cognitive load because the visual input is static: only a fixation cross remains on the screen during presentation of the digits and the retention interval. Therefore, neither pupil size nor EEG alpha activity are affected by visual input, but only by changes in the cognitive load. As compared to most common modern EEG WM paradigms, we presented the items not simultaneously but sequentially every two seconds making it possible to study neural activity during encoding and maintenance of each presented memory item at a fine time scale. Moreover, we introduced a control condition with passive listening to differentiate pure WM processes from auditory perceptual processes.

The richness and simplicity of the dataset opens an opportunity to researchers of individual differences, especially those who are interested in how the human brain deals with cognitive overload, as well as computational neuroscientists and engineers looking for a testing bed of cognitive load classification algorithms.

The dataset has never been used in any publications.

## Methods

### Participants

Eighty-six participants completed the task. The participants had normal or corrected-to-normal vision and reported no history of neurological or mental diseases. All the participants were Russian native speakers. None of the participants had had any disease of the nervous system, psychiatric, or hearing disorders in the past, or reported use of any medication. Informed consent was obtained from each participant. The experimental protocol was approved by the Ural Federal University ethics committee.

A general description of the sample is provided in Table 1. We used the Annett Handedness Scale (Annett, 1970) to determine the handedness. Ocular dominance was determined with three different tests: Rosenbach’s test (Rosenbach, 1903), aiming (the participants were asked to make the finger gun gesture with both hands and then aim to a self-selected target while closing one of the eyes (the other one is the dominant eye)), and hole-in-the-card (Fink, 1938).

**Table 1.** Participant overview

|  |  |
| --- | --- |
| **Handedness** | 3 ambidextrous, 6 left-, 77 right-handed |
| **Dominant eye** | 25 left, 61 right |
| **Sex** | 74 females, 12 males |
| **Age (mean ± SD)** | 20.5 ± 3.9 years |

### Task and procedure

Before the WM task, we recorded 4 minutes of resting state EEG. The participants were seated in a comfortable chair and asked to close their eyes and to sit still. After the resting state recording, the participants were given instructions and proceeded with the WM task (see Figure 1).

**Figure 1.**
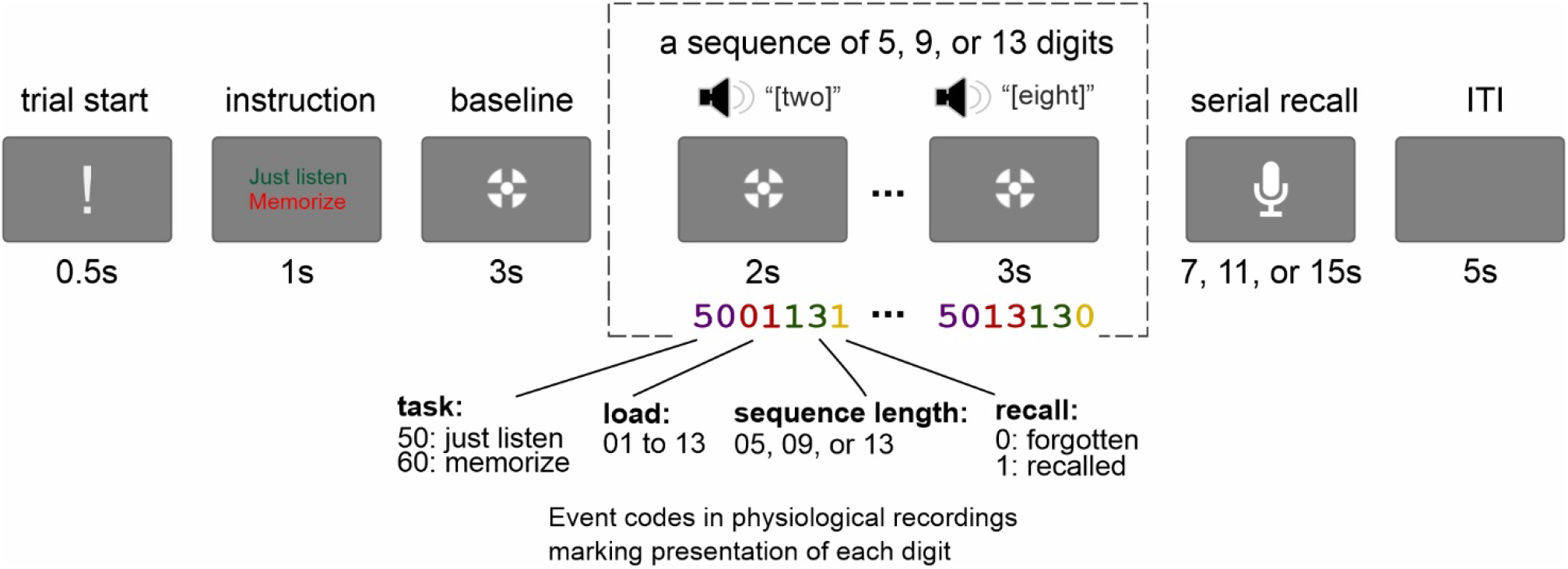
A visual depiction of the experimental paradigm (not in scale). The participants were instructed to either memorize or just listen to sequences of 5, 9, or 13 digits, and then in the memory condition, they recalled the whole sequence in the serial order. The presentation of each digit is marked in the physiological recordings with event codes varying from 500105 to 6013131. For example, event code 5002091 means that the task was to passively listen to the sequence, the digit is the second in a sequence of nine digits, and the digit was correctly recalled.

Each trial began with an exclamation mark for 0.5 s along with a recorded voice command “begin” – indicating the start of the trial. The exclamation mark was followed by an instruction to either memorize the subsequent digits in the correct order (memory condition) or to just listen to the digits without attempting to memorize them (control condition). The instruction was followed by a three-second baseline period. Then either 5, 9, or 13 digits were presented auditorily with an SOA of 2 seconds. The digits were presented with a female voice in Russian. Each of the digits from 0 to 9 was used, and the mean duration of each digit was 664 ms (min: 462 ms, max: 813 ms). The last digit in the sequence was followed by a 3-sec retention interval. During the baseline, encoding, and retention intervals, participants were fixating a cross (1.2 cm in diameter) on the screen. In the memory condition, the participants were asked to recall each digit out loud in the correct order starting from the first one (i.e., serial recall). The retrieval was recorded by a computer microphone controlled by PsychoPy (Peirce, 2007). The participants had 7, 11, and 15 seconds for 5, 9, and 13 digit sequences, respectively, to recall the digits. The retrieval was followed by an inter-trial interval (ITI) of 5 s. In the control condition (passive listening), presentation of the digits and the delay period was followed immediately by an ITI of the same duration.

There were 9 blocks in total with 54 passive listening and 108 memory trials overall. Each block consisted of 3 control (one of each load) followed by 12 memory (4 trials on each level of load, in random order) followed again by 3 control trials. Before the main WM task, each participant completed 6 practice trials (3 passive listening and 3 memory trials). After each block, the participant had a self-paced break. After every 3 blocks, the participants took a longer break when they could take a snack and filled out a NASA-TLX questionnaire to self-assess the level of mental, physical, and temporal demand they experienced, their perceived overall performance, effort, and frustration during the preceding three blocks of the task.

The presentation script written in PsychoPy is available in the “stimuli” folder of the database.

### Recording setup

A 64-channels EEG system with active electrodes (ActiCHamp, Brain Products, Germany) was used for the recording. The electrodes were placed according to the extended 10-20 system with FCz channel as the online reference and Fpz as the ground electrode. The level of impedance was maintained below 25 kOm. The sampling rate was 1000 Hz. No online digital filters were applied.

Cardiovascular measures (ECG and PPG) were acquired using the same amplifier from the auxiliary (AUX) inputs. ECG was recorded from one channel with the active electrode placed on the right wrist and the reference electrode on the left wrist, and the ground on the left inner forearm at 3 cm distally from the elbow. Finger photoplethysmography was recorded from the left index finger.

Pupillometry was recorded with Pupil Labs wearable eye-tracker (Pupil Lab GmBH, Germany) with 120 Hz sampling rate. One-point calibration preceded each recording.

The distance to the monitor was 80 cm. Two loudspeakers were placed on the sides of the monitor with 84 cm between them (62°). The measured loudness of the digit sounds was 70 dB SPL. The loudness was measured by placing a Mastech MS6701 sound level meter near where the participants’ head was located. The luminance in the room was set to 380 lux.

### Data Records

The data are uploaded to OpenNeuro (https://openneuro.org, https://openneuro.org/datasets/ds003838) in BIDS format (Pernet et al., 2019) (see Figure 2 for the data structure). “participants.tsv” spreadsheet in the main folder contains information about the participants such as age, sex, handedness, dominant eye, and NASA-TLX scale results. More detailed description of the variables is available in the corresponding “.json” file. The folder “stimuli” in the main folder contains the presentation script written in PsychoPy and a graphical representation of the task with the corresponding event codes (same as Figure 1 here). Other 86 folders, one per participant, contain the data: “eeg”– raw EEG data, “ecg”– raw ECG and PPG data, “beh”– behavioral data, “pupil”– raw pupillometry and eye-tracking data. Pupillometry data are saved in “.tsv” format that combines original “pupil_positions.csv” and “gaze_positions.csv” raw data files and identified blinks exported from the Pupil Player software. “eeg” and “ecg” types of data saved in the EEGLAB file format (.set files). The “eeg” and “ecg” data are available during resting state with eyes closed (“rest” task) and during the working memory task (“memory” task). There is no resting state data for pupillometry. The “beh” type of data are shared as spreadsheets including participants’ trial-by-trial and item-by-item responses in the working memory task (only memory condition). The behavioral responses were manually transcribed from the recorded speech by at least two scorers and the mismatching trials were then checked by one of the coauthors. The codebooks for each type of data for events and variables are available in each folder in the corresponding “.json” files.

**Figure 2.**
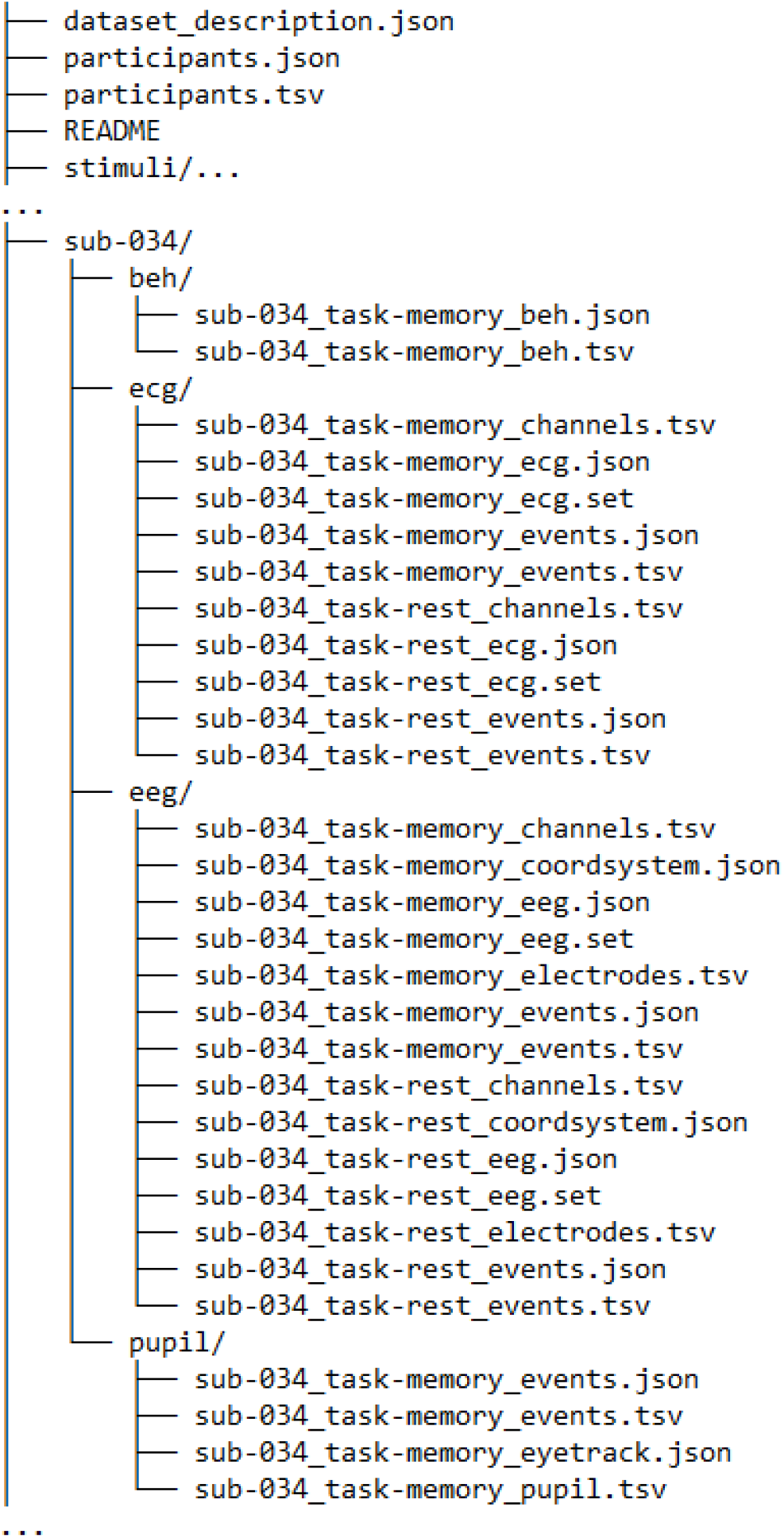
Data structure

Not all data are available in every data modality. Three EEG and two pupillometry datasets were excluded because of technical failure. 19 EEG recordings were excluded due to an experimenter error resulting in the misplaced electrode cap being incompatible with 10-20 system electrode layout. Thus, this dataset consists of 65 EEG, 83 ECG+PPG, and 84 pupillometry recordings. Behavioral data are available for all 86 participants. The missing data in each modality is labeled in the “participants.tsv” file.

### Technical Validation

#### Behavior

The average number of recalled digits in the correct order was 4.62 ± 0.75, 4.18 ± 0.56, 3.57 ± 0.55 for 5-, 9-, and 13-digit sequences, respectively. See Figure 3a for a visual depiction.

**Figure 3.**
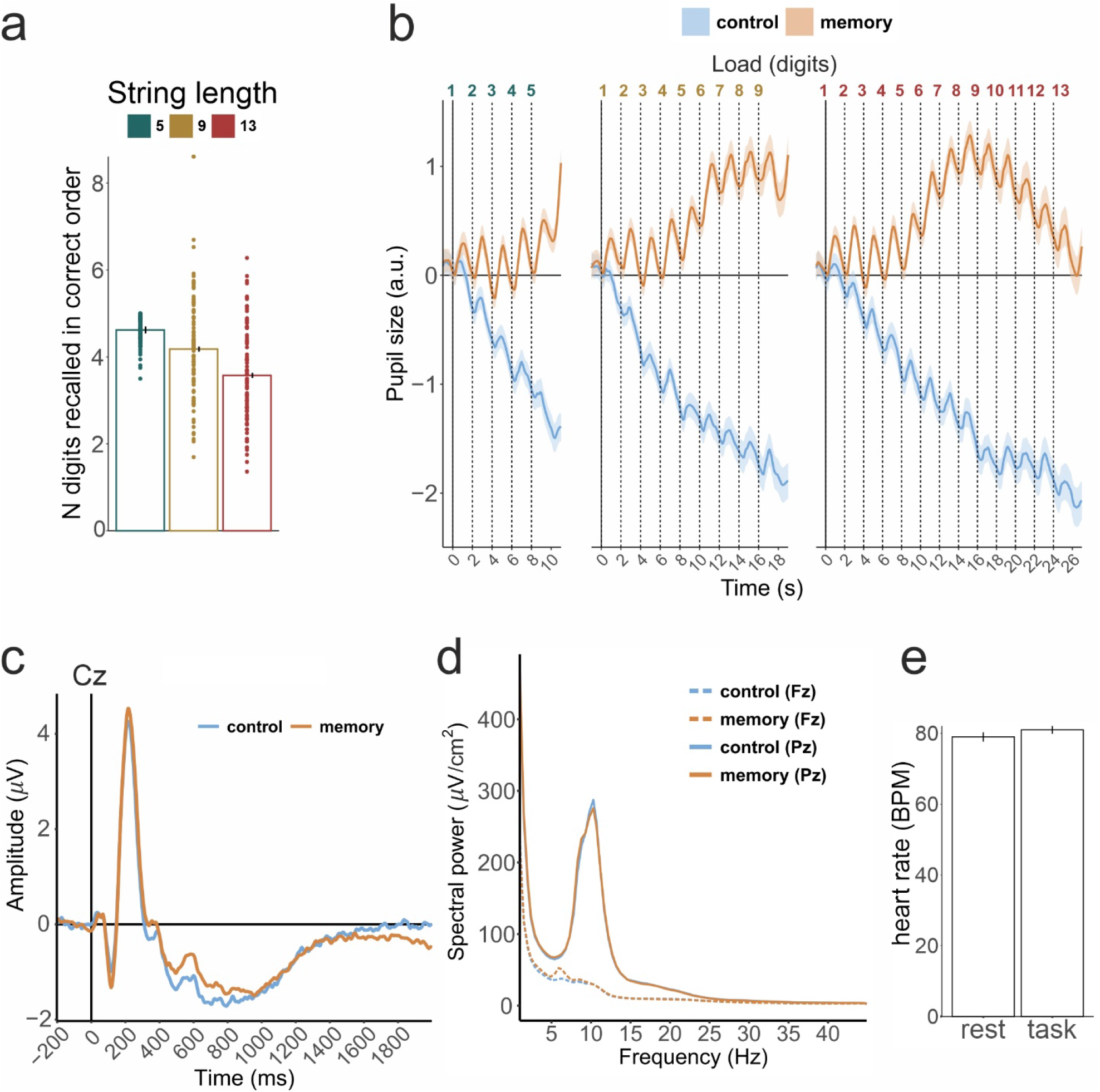
Validation analysis results. (a) Behavioral performance with individual participant data points for the three sequence lengths. Error bars are the standard errors of the mean. (b) Pupillometry data averaged over 5-, 9-, and 13-digit sequence lengths (left, middle, and right panels, respectively) in two conditions. Vertical dashed lines represent presentation of the digits. Shading is the standard error of the mean. (c) Event-related potentials with average mastoid reference at Cz channel averaged over all digits and sequence lengths in passive listening (control) and memorizing (memory) conditions. (d) Absolute spectral power after current source density (CSD) transformation at Fz and Pz channels averaged over 2 second epochs corresponding to encoding and maintenance of single digits in all conditions. For ERP and spectral power analyses, the artifacts were first suppressed by means of independent component analysis (ICA) and then visually identified epochs still containing artifacts were rejected. (e) Heart rate in beats per minute in the resting state and during the tasks as derived from ECG. Error bars are the standard errors of the mean.

#### Pupillometry

For the validation analysis, the pupillometry data were preprocessed with gazeR package in R (Geller et al., 2019), and baseline corrected by subtracting the mean absolute value in the interval of 2 seconds before presentation of the first digit in the sequence. To assure that our strongest experimental manipulation elicited expected changes in pupil size, we averaged the data in the control and memory conditions. The pupil size showed both phasic (with two seconds after stimulus presentation) and tonic responses throughout the trial (see Figure 3b). Similar to previous studies, we observed an increase in pupil size in the active memory condition, and a sustained downward trend during passive listening (Granholm et al., 1996; Peavler, 1974).

#### EEG

We first visually inspected EEG traces and assured absence of large potentially unusable or missing chunks of data. Additionally, we performed two other checks of the data quality. In an analysis of the absolute spectral power, we observed the typical alpha peak at around 10 Hz at posterior channels, which did not differ between the memory and control condition. Furthermore, confirming expectations from verbal WM EEG literature (Pavlov & Kotchoubey, 2020, in press), we observed an increase in frontal midline theta activity at around 6 Hz in the memory condition compared to the control condition (see Figure 3d). Event-related potentials in response to the verbal material demonstrated clearly visible early N1-P2 components with near zero baseline amplitude (Figure 3c).

#### ECG and PPG

The ECG and PPG traces were visually inspected to detect the presence of R peaks and pulse waves for most of the recording time. Extracted from both channels, the average heart rate was almost exactly the same in ECG and PPG data, and fell within the normal range in the resting state (79 ± 12 bpm) and the task (81 ± 10 bpm) (Figure 3e).

## Code availability

The presentation script is shared within the BIDS formatted database.

## Author contributions statement

Yuri G. Pavlov: Conceptualization, Funding acquisition, Data curation, Formal analysis, Project administration, Supervision, Visualization, Methodology, Writing - original draft, Writing - review & editing.

Alexander I. Kotyusov: Investigation, Methodology, Data curation.

Dauren Kasanov: Investigation, Methodology, Data curation.

Alexandra I. Kosachenko: Investigation, Methodology, Data curation, Writing - review & editing.

Niko A. Busch: Validation, Writing - review & editing.

## Potential Conflicts of Interest

Nothing to report.

## Acknowledgements

This study was supported by Russian Foundation for Basic Research (RFBR) #19-013-00027. Open access publication was partially supported by German Research Society (Deutsche Forschungsgemeinschaft, DFG), grant KO-1753/13-4.

## Notes

https://openneuro.org/datasets/ds003838

### Competing Interest Statement

The authors have declared no competing interest.

## References

Annett, M. (1970). A Classification of Hand Preference by Association Analysis. British Journal of Psychology, 61(3), 303–321. https://doi.org/10.1111/j.2044-8295.1970.tb01248.x

Constantinidis, C., & Klingberg, T. (2016). The neuroscience of working memory capacity and training. Nature Reviews Neuroscience, 17(7), 438–449. https://doi.org/10.1038/nrn.2016.43

Fink, W. H. (1938). The dominant eye: Its clinical significance. Archives of Ophthalmology, 19(4), 555–582. https://doi.org/10.1001/archopht.1938.00850160081005

Fukuda, K., Vogel, E., Mayr, U., & Awh, E. (2010). Quantity, not quality: The relationship between fluid intelligence and working memory capacity. Psychonomic Bulletin & Review, 17(5), 673–679. https://doi.org/10.3758/17.5.673

Geller, J., Winn, M., Mahr, T., & Mirman, D. (2019). GazeR: A package for processing gaze position and pupil size data. PsyArXiv. April, 22.

Granholm, E., Asarnow, R. F., Sarkin, A. J., & Dykes, K. L. (1996). Pupillary responses index cognitive resource limitations. Psychophysiology, 33(4), 457–461.

Jacobs, J. (1887). Experiments on” prehension”. Mind, 12(45), 75–79. https://doi.org/10.1093/mind/os-12.45.75

Johnson, M. K., McMahon, R. P., Robinson, B. M., Harvey, A. N., Hahn, B., Leonard, C. J., Luck, S. J., & Gold, J. M. (2013). The relationship between working memory capacity and broad measures of cognitive ability in healthy adults and people with schizophrenia. Neuropsychology, 27(2), 220–229. https://doi.org/10.1037/a0032060

Joshi, S., Li, Y., Kalwani, R. M., & Gold, J. I. (2016). Relationships between Pupil Diameter and Neuronal Activity in the Locus Coeruleus, Colliculi, and Cingulate Cortex. Neuron, 39(1), 221–234. https://doi.org/10.1016/j.neuron.2015.11.028

Luck, S. J., & Vogel, E. K. (2013). Visual working memory capacity: From psychophysics and neurobiology to individual differences. Trends in Cognitive Sciences, 17(8), 391–400. https://doi.org/10.1016/j.tics.2013.06.006

Murphy, P. R., O’Connell, R. G., O’Sullivan, M., Robertson, I. H., & Balsters, J. H. (2014). Pupil diameter covaries with BOLD activity in human locus coeruleus. Human Brain Mapping, 35(8), 4140–4154. https://doi.org/10.1002/hbm.22466

Pavlov, Y. G., & Kotchoubey, B. (2020). The electrophysiological underpinnings of variation in verbal working memory capacity. Scientific Reports, 10(1), 16090. https://doi.org/10.1038/s41598-020-72940-5

Pavlov, Y. G., & Kotchoubey, B. (in press). Oscillatory brain activity and maintenance of verbal and visual working memory: A systematic review. Psychophysiology, e13735. https://doi.org/10.1111/psyp.13735

Peavler, W. S. (1974). Pupil size, information overload, and performance differences. Psychophysiology, 11(5), 559–566.

Peirce, J. W. (2007). PsychoPy—Psychophysics software in Python. Journal of Neuroscience Methods, 162(1), 8–13. https://doi.org/10.1016/j.jneumeth.2006.11.017

Pernet, C. R., Appelhoff, S., Gorgolewski, K. J., Flandin, G., Phillips, C., Delorme, A., & Oostenveld, R. (2019). EEG-BIDS, an extension to the brain imaging data structure for electroencephalography. Scientific Data, 6(1), 103. https://doi.org/10.1038/s41597-019-0104-8

Rosenbach, O. (1903). Ueber monokulare Vorherrschaft beim binokularen Sehen. Muenchener Medizinische Wochenschrift, 30, 1290–1292.

